# Transformers Outperform ConvNets for Root Segmentation: A Systematic Comparison Across Nine Datasets

**DOI:** 10.64898/2026.02.18.706562

**Authors:** Abraham George Smith, Sotiris Lamprinidis, Anand Seethepalli, Larry M. York, Eusun Han, Patrick Möhl, Kyriaki Boulata, Kristian Thorup-Kristensen, Jens Petersen

## Abstract

Root segmentation is a fundamental yet challenging task in image-based plant phenotyping. We present the first systematic comparison of Transformer and Convolutional Neural Network (ConvNet) architectures for root segmentation, evaluating 21 architectures across nine diverse datasets and comparing pre-trained models to training from scratch. Transformer-based models significantly outperform ConvNets for segmentation accuracy and root-diameter agreement. Pre-training significantly improves mean Dice from 0.623 to 0.666 (*p* = 3.3 × 10^−10^). We also find that Transformers benefit more from pre-training than ConvNets, with Dice improvements of +0.072 versus +0.022 (*p* = 3.7 × 10^−4^), supporting the hypothesis that fine-tuned Transformers transfer more effectively across large domain gaps. Among evaluated models, MobileSAM achieved the highest Dice score while maintaining computational efficiency. Dataset choice explained far more performance variance (70.9%) than model architecture (6.7%), suggesting that data curation matters more than model selection.

## Introduction

Image-based methods are increasingly used to quantify plant root traits [Weihs et al., 2024]. Segmentation of images into roots and background is a prerequisite for the extraction of root traits relevant to plant physiology [Möhl et al., 2022], breeding [Kuijken et al., 2015], and agronomy [Thorup-Kristensen et al., 2020a].

Smith et al. [2020] demonstrated U-Net’s [Ronneberger et al., 2015] efficacy for the segmentation of chicory root images in soil. Their solution used images from a controlled Rhizobox setup [Thorup-Kristensen et al., 2020b].

More recently, complex root datasets presenting variation and artifacts not yet solved by U-Net [Handy et al., 2024] have highlighted that fully automatic root segmentation remains challenging. Even for less challenging Rhizotron datasets, treatment effects or genotype differences may be subtle and only revealed by sufficiently accurate segmentation models. Baykalov et al. [2023]compared segmentation methods across diverse rhizotron datasets, finding that U-Net architectures achieved the best accuracy but that heterogeneous datasets remained challenging.

Transformer-based architectures have recently been applied to root segmentation. Li et al. [2025] showed that Swin-Unet++ outperformed U-Net by 1.08% mIoU on cabbage seedling images, and Zhou et al. [2024] applied Point Transformers to 3D Arabidopsis root point clouds. For ConvNet comparisons, Shen et al. [2020] found that an improved DeepLabv3+ outperformed U-Net for cotton root segmentation from Minirhizotron images. However, these studies each evaluated one or two architectures on a single dataset. No systematic comparison of multiple Transformer and ConvNet architectures has been conducted across diverse root imaging conditions, nor have vision foundation models such as SAM been evaluated for root segmentation.

### Contributions

1. The first systematic comparison of Transformer and ConvNet architectures for root segmentation across diverse datasets
2. Empirical evidence that Transformers benefit more from pre-training than ConvNets when the domain gap is large
3. Practical recommendations for model selection in root phenotyping pipelines

We hypothesise that: (H1) Transformer architectures outperform ConvNets for root segmentation, and (H2) pre-training improves performance for both architecture families compared to training from scratch.

To test these hypotheses, we evaluate 21 architectures across nine root image datasets, comparing pre-trained models to models trained from scratch. We include vision foundation models (MobileSAM, SAM2) and recent ConvNets (U-Net++, DeepLabV3+, MAnet). In total, we trained 1,511 models to assess all combinations of architecture, dataset, pre-training, and learning rate, producing over 3 million segmentations for evaluation.

## Methods

### Datasets

We evaluated model performance on nine publicly available datasets spanning diverse species, root appearances, and imaging conditions. All datasets are open access, supporting reproducibility. We selected these datasets to represent variation in imaging modality, scale, and annotation density. The datasets are described below with references to the original publications. Example images are shown in Figure 2; dataset splits, image resolutions, and DPI are provided in Supplementary Table 3.

**Figure 1.**
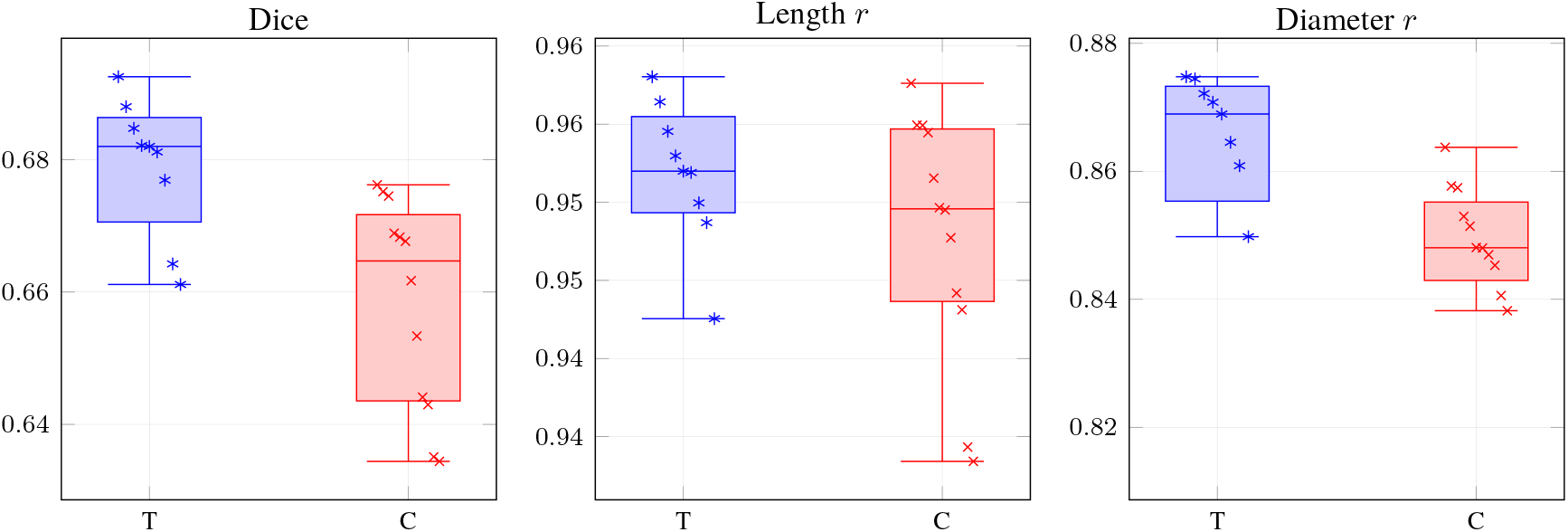
Per-model test performance by architecture family. T = Transformer, C = ConvNet. Left: Dice. Middle: Length *r*. Right: Diameter *r. r* = Pearson correlation coefficient. (One value per model; configuration selected by mean validation Dice across datasets).

**Figure 2.**
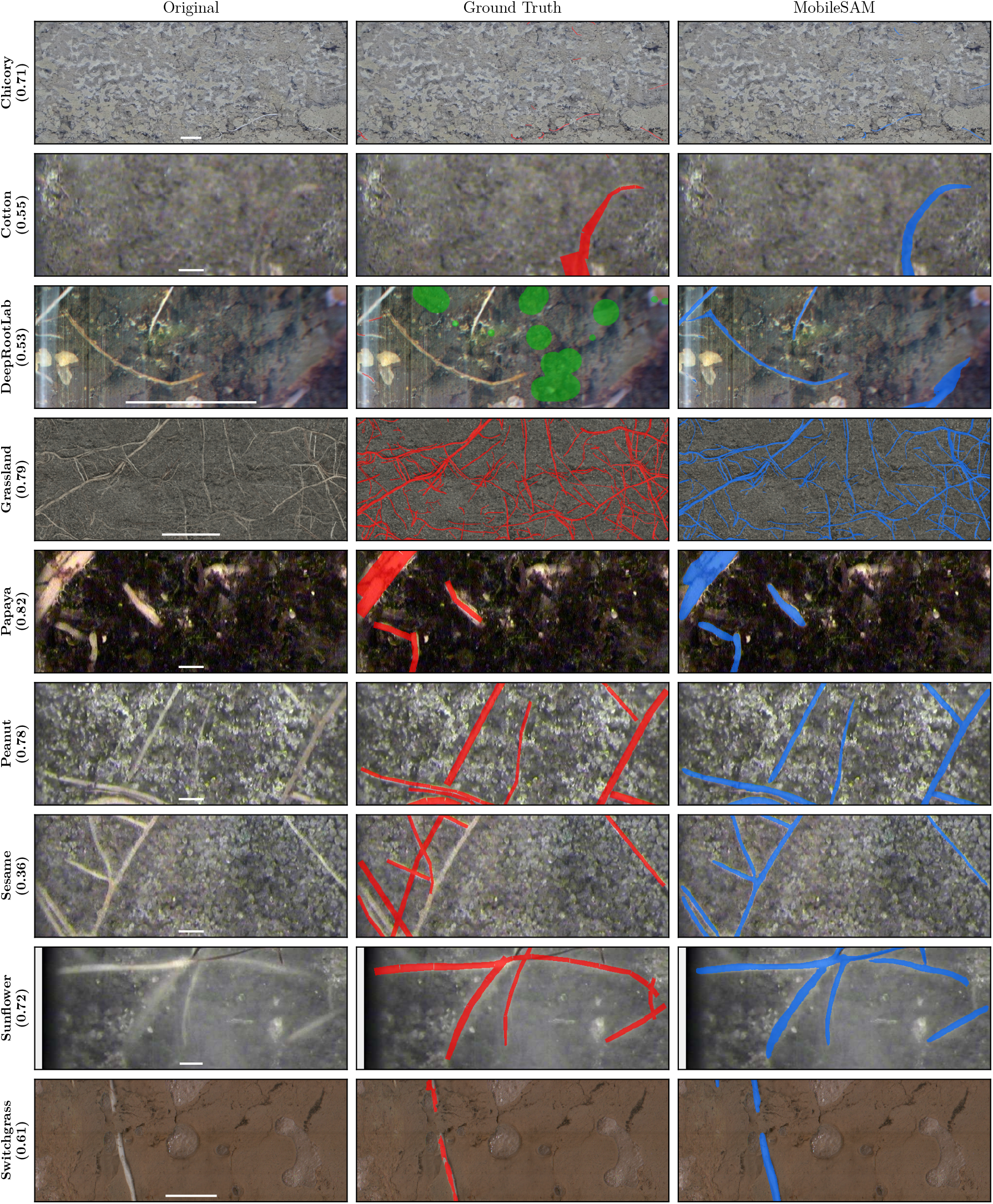
Qualitative comparison of MobileSAM predictions. Each row shows the test image with median Dice for that dataset: original image (left), ground truth overlay in red (middle), and MobileSAM prediction overlay in blue (right). Chicory is cropped to the central 25% for visibility. Scale bars (1 cm) shown in bottom center of original images.

#### DeepRootLab

This dataset includes images of eleven species, collected and annotated as part of the Five Seasons project [Han et al., 2025]. The species are: intermediate wheatgrass (*Thinopyrum intermedium*), perennial lupine (*Lupinus perennis*), chicory (*Cichorium intybus*), dyers woad (*Istatis tinctoria*), mugwort (*Artemisia vulgaris*), rosinweed (*Silphium perfoliatum*), comfrey (*Symphytum officinale*), curly dock (*Rumex crispus*), tall fescue (*Festuca arundinacea*), winter rye (*Secale cereale*), and winter wheat (*Triticum aestivum*). Annotations represent corrective edits rather than complete root traces; consequently, this dataset is excluded from root-length and root-diameter correlation analyses. Images are available from https://zenodo.org/records/15213661.

#### Grassland

Collected in alpine grasslands and released as part of a study on root growth and soil function [Möhl et al., 2022], this dataset captures natural root-soil interactions under field conditions. Images are available from https://figshare.com/ndownloader/articles/20440497/versions/2.

#### Chicory

We used the densely annotated subset of the chicory dataset introduced in [Smith et al., 2020] and published in [Smith et al., 2019]. Images are available from https://zenodo.org/records/3527713.

#### PRMI Collection

These six datasets, papaya (*Carica papaya*), peanut (*Arachis hypogaea*), sesame (*Sesamum indicum*), sunflower (*Helianthus annuus*), cotton (*Gossypium hirsutum*), and switchgrass (*Panicum virgatum*), were released as part of the PRMI benchmark for minirhizotron root imaging [Xu et al., 2022]. Each dataset corresponds to a different crop species and imaging context. Images are available from https://gatorsense.github.io/PRMI/.

We used the original dataset splits where available (PRMI benchmark, Grassland, Chicory). For DeepRootLab, we split images 60/20/20 into training, validation, and test sets.

### Neural network architectures

The evaluated architectures (Table 1) include both convolutional and Transformer models, with and without pre-trained weights.

**Table 1.** Evaluated segmentation architectures, encoders, and pre-training sources. ^a^ Excluded from pre-trained evaluation on Chicory to prevent data leakage.

| Architecture | Encoder | Pre-training | Type |
| --- | --- | --- | --- |
| U-Net++ | ResNet50 [He et al., 2016]<br>InceptionV4 [Szegedy et al., 2017] | ImageNet [Russakovsky et al., 2015] | ConvNet |
| LinkNet | ResNet50<br>InceptionV4 | ImageNet | ConvNet |
| MAnet | ResNet50<br>InceptionV4 | ImageNet | ConvNet |
| SegRoot | w8d5 [Wang et al., 2019] | SegRoot | ConvNet |
| Mask2Former | ResNet50<br>Swin-Tiny<br>Swin-Small [Liu et al., 2021b] | COCO [Lin et al., 2014] | Transformer |
| SegFormer | mit-b1<br>mit-b2<br>mit-b3 [Xie et al., 2021] | Cityscapes [Cordts et al., 2016] | Transformer |
| UNetGN | GroupNorm [Smith et al., 2020] | Chicory<sup>a</sup> | ConvNet |
| UNetGNRes | GroupNorm + res. [Smith et al., 2022] | Chicory<sup>a</sup> | ConvNet |
| DeepLabV3/V3+ | DeepLabV3 (ResNet50)<br>DeepLabV3+ (ResNet50) | ImageNet | ConvNet |
| RootNav 2.0 | Stacked hourglass [Newell et al., 2016] | RootNav | ConvNet |
| MobileSAM | ViT-Tiny [Dosovitskiy et al., 2021] | SAM [Kirillov et al., 2023] | Transformer |
| SAM2 | Hiera-Small<br>Hiera-Base+ | SAM2.1 [Ravi et al., 2024] | Transformer |

### Convolutional Architectures

These include encoder-decoder models widely used in biomedical and plant image analysis, as well as several lightweight or attention-augmented variants:

- **UNetGN** and **UNetGNRes**, both based on the original U-Net architecture [Ronneberger et al., 2015], were adapted for root segmentation by replacing BatchNorm with GroupNorm to better accommodate small batch sizes. UNetGN was introduced in [Smith et al., 2020], while UNetGNRes is a residual variant with reduced parameters, developed as part of Root-Painter [Smith et al., 2022].
- **U-Net++** [Zhou et al., 2018], which enhances the U-Net architecture with nested and dense skip connections. We used the implementation from Segmentation Models Pytorch [Iakubovskii, 2019].
- **DeepLabV3** and **DeepLabV3+** [Chen et al., 2017, 2018], encoder-decoder models using dilated convolutions and ResNet-50 backbones. The implementation for DeepLabV3 was taken from TorchVision [Paszke et al., 2019], while the DeepLabV3+ implementation was from Segmentation Models Pytorch [Iakubovskii, 2019].
- **LinkNet** [Chaurasia and Culurciello, 2017] and **MAnet** [Fan et al., 2020], offering residual or attention-based improvements over basic encoder-decoder structures. The implementations were taken from Segmentation Models Pytorch [Iakubovskii, 2019].
- **RootNav 2.0** [Yasrab et al., 2019], a deep hourglass-style network for root system segmentation.
- **SegRoot** [Wang et al., 2019], a compact convolutional architecture tailored for fine root segmentation.

### Transformer Architectures

These models incorporate self-attention mechanisms and hierarchical token processing for segmentation:

- **SegFormer** [Xie et al., 2021], a lightweight transformer architecture with multiscale hierarchical encoding and efficient MLP decoding. We used the implementation from Segmentation Models Pytorch [Iakubovskii, 2019], evaluated using MIT-B1, MIT-B2 and MIT-B3 backbones.
- **Mask2Former** [Cheng et al., 2021], a unified architecture for panoptic and semantic segmentation, tested with ResNet-50, Swin-Tiny, and Swin-Small backbones.
- **MobileSAM** [Zhang et al., 2023], a mobile-optimized adaptation of the Segment Anything model, using a ViT-Tiny backbone.
- **SAM2** [Ravi et al., 2024], the second generation Segment Anything model for images and video, tested with Hiera-Small and Hiera-Base+ backbones.

### Pre-training

Models were evaluated with pre-trained weights and trained from scratch with random initialisation. Pre-training sources are listed in Table 1.

### Training procedure

Each model was trained with two learning rates (0.001 and 0.0001), both with pre-trained weights and from scratch. Each of these four configurations was trained twice with different random seeds, yielding 8 training runs per model per dataset. Across 21 models and 9 datasets, this resulted in 1,511 training runs. One run failed to converge (MobileSAM ViT-T on DeepRootLab, from scratch at a learning rate of 0.001), never exceeding 0.0 validation Dice.

Training used the AdamW optimiser with 16-bit mixed precision. Each run was limited to a maximum of 2 hours. A warm-up schedule of 2 epochs was applied, followed by early stopping if validation Dice did not improve for 20 epochs. The loss function combined Dice loss and cross-entropy, which has proven effective for class-imbalanced root datasets [Smith et al., 2020]. To mitigate class imbalance, batches containing no root annotations were excluded.

Patch sizes were selected based on architecture requirements: 572 pixels for UNet-GN, 576 pixels for models requiring dimensions divisible by 64, and 1024 pixels for transformer-based architectures. Batch sizes were set to the largest value in { 4, 8, 16} that fit in GPU memory.

Data augmentation was applied on the fly during training and included colour jitter (66% probability), greyscale conversion (5%), pixel inversion (2.5%), random rotation ( ± 15^°^, 33%), perspective distortion (33%), horizontal flipping (50%), and elastic deformation (*α*=75, 33%).

All training was performed on two machines with identical hardware, each equipped with an AMD Ryzen 9 7950X CPU, 32GB RAM, and an NVIDIA RTX 4090 GPU with 24GB VRAM. Computational cost was quantified in billions of floating point operations (GFLOPs) per forward pass, computed using the calflops library [MrYxJ, 2023], and number of parameters, as these affect hardware requirements, running costs, and environmental impact. To jointly compare efficiency and accuracy, we ranked models by the mean of their Dice, parameter count, and FLOPs ranks, providing a simple combined metric for practitioners balancing these trade-offs. Training code is available at https://github.com/sotlampr/seg.

### Configuration selection

To prevent overfitting to the test set, model selection used a two-stage procedure based on validation performance:

### Replicate selection

For each combination of model, dataset, learning rate, and pre-training, the replicate with the highest validation Dice was retained, along with its paired test result.

### Hyperparameter selection

For each model, the configuration (learning rate, pre-training) with the highest mean validation Dice across all datasets was selected.

All reported test metrics use the configuration selected by this procedure.

For pre-trained vs scratch comparisons, configuration selection was performed separately within each condition: for each model, the best learning rate was selected independently for pre-trained runs and for scratch runs, both based on mean validation Dice across datasets. This gave paired observations for each model, one pre-trained and one trained from scratch.

### Trait extraction

Root morphological traits were extracted from both predicted segmentations and ground-truth annotations using a fork of RhizoVision Explorer (RVE) [Seethepalli et al., 2021] with bug fixes and headless functionality (see Supplementary Materials), parallelized with GNU Parallel [Tange, 2018]. For each segmentation mask, we computed total root length and mean root diameter. These traits allow evaluation beyond pixel-level accuracy, assessing whether models produce segmentations that yield biologically meaningful measurements. Total root length is relevant for rooting density and specific root length, a widely measured functional trait, while mean diameter captures root thickness which relates to root development and function.

Additionally, we configured RVE to compute root length binned by diameter in 1-pixel increments, enabling analysis of how root length is distributed across different root diameters. This diameter distribution reveals whether models accurately capture the full range of root sizes, particularly thin roots which are challenging to segment.

### Evaluation metrics

#### Dice coefficient

Primary segmentation quality metric, computed as the harmonic mean of precision and recall between predicted and ground-truth masks.

#### Root-length correlation

Pearson correlation coefficient between predicted and ground-truth total root length, measuring agreement in overall root quantity estimation.

#### Root-diameter correlation

Pearson correlation coefficient between predicted and ground-truth mean root diameter, measuring agreement in root thickness estimation.

### Statistical analysis

To compare Transformer and ConvNet architectures (a between-model comparison), we averaged each model’s metric across datasets and applied an independent two-sample *t*-test on model means (*n* = 9 Transformer, *n* = 12 ConvNet) with a one-sided alternative (Transformer *>* ConvNet). For pre-training comparisons and the architecture-by-pre-training interaction (within-model contrasts across datasets), we used linear mixed effects models with dataset and model as random intercepts to account for the repeated-measures structure. The pre-training comparison used a one-sided alternative (pre-trained *>* scratch); the interaction test was two-sided.

All tests used a significance level of *α* = 0.05. All configuration selection was performed on validation data; all reported metrics are from held-out test sets.

### Thin root discrepancy analysis

To understand the sources of thin root discrepancy between model predictions and annotations, we selected the 10 images with the largest root length underestimation per diameter bin (bins 1–6, in 1-pixel increments) for MobileSAM across all eight datasets, or fewer where a bin contained fewer than 10 images, yielding 437 images. Each image was visually inspected and the dominant source of error was identified: an RVE measurement artifact (where annotation corners were falsely detected as thin roots), roots missed entirely by the model, or roots predicted too thick (model segmentation wider than the annotation). Qualitative examples (Figures 6 and 7) were selected as the images with the largest discrepancy for each error type.

## Results

### Transformers outperform ConvNets

Transformer models had significantly higher mean test Dice than ConvNets (0.679 vs 0.659; *p* = 1.5 × 10^−3^; Table 2). This difference is visible in Figure 1 (left). MobileSAM ViT-T, M2F Swin-S, and M2F Swin-T had the highest individual Dice values, occupying the top three positions in the ranking (Table 5). Several ConvNets still reached Dice values comparable to mid-ranking Transformers, including MA-Net Inc-v4 and U-Net++ Inc-v4. Mean test root-length correlation was higher for Transformers than ConvNets (0.952 vs 0.948; *p* = 0.099; Table 2), though this difference was not statistically significant. Overlap is visible in the per-model distributions (Figure 1, middle). SegFormer B1 had the highest root-length correlation (Supplementary Table 1).

**Table 2.**
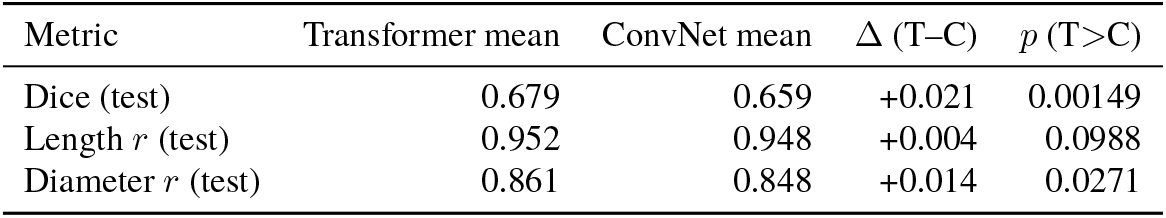
Summary of architecture-family differences (Transformers vs ConvNets) across metrics.

| Metric | Transformer mean | ConvNet mean | $\Delta$ (T-C) | $p$ (T>C) |
| --- | --- | --- | --- | --- |
| Dice (test) | 0.679 | 0.659 | +0.021 | 0.00149 |
| Length $r$ (test) | 0.952 | 0.948 | +0.004 | 0.0988 |
| Diameter $r$ (test) | 0.861 | 0.848 | +0.014 | 0.0271 |

**Table 3.**
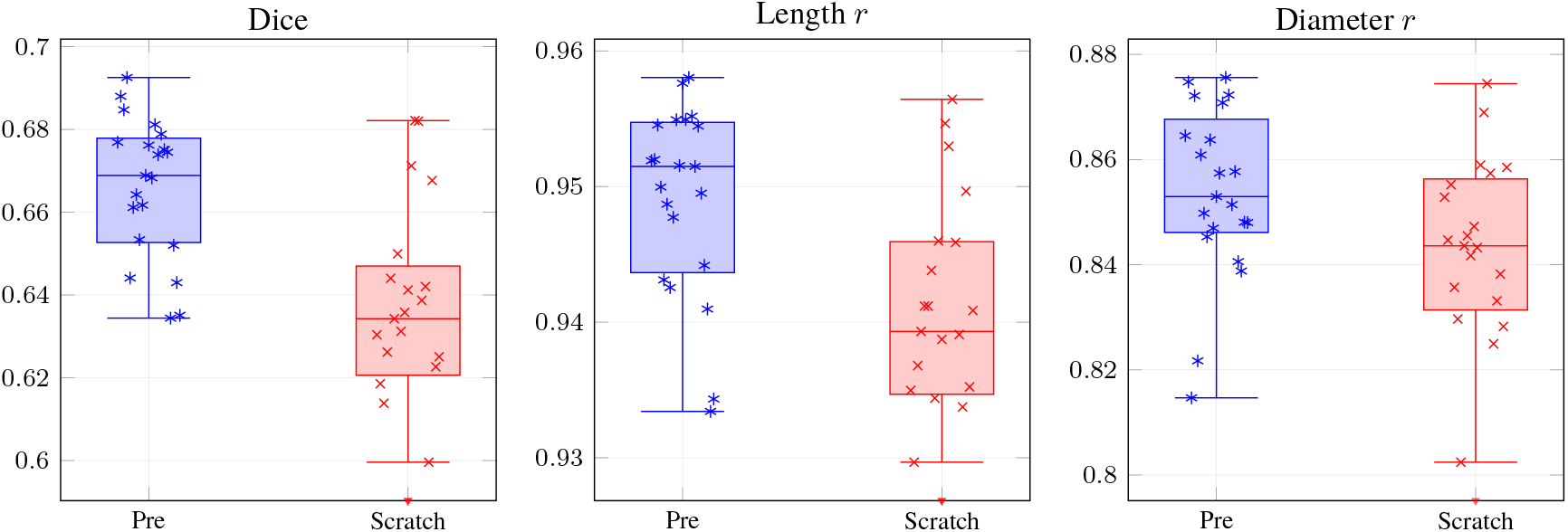
Per-model test performance by pretraining. Left: Dice. Middle: Length *r*. Right: Diameter *r. r* = Pearson correlation coefficient. Scratch-trained SAM2 models excluded as outliers.

Mean test root-diameter correlation was significantly higher for Transformers than ConvNets (0.861 vs 0.848; *p* = 0.027; Table 2). The top-ranked models for diameter correlation were predominantly Transformers: Seg-Former B2, M2F Swin-S, and SegFormer B3 (Supplementary Table 2).

### Pre-training improves performance

Pre-trained models had significantly higher mean test Dice than models trained from scratch (0.666 vs 0.623; *p* = 3.3 × 10^−10^; Table 4; Figure 3). Pre-trained models also had significantly higher root-length correlation (0.949 vs 0.907; *p* = 1.6 × 10^−4^) and root-diameter correlation (0.854 vs 0.828; *p* = 2.3 × 10^−3^). The highest-performing models, including MobileSAM ViT-T, M2F Swin-S, and SegFormer B2, were pre-trained, while the lowest-performing were trained from scratch (Table 5; Supplementary Tables 1 –2).

**Table 4.**
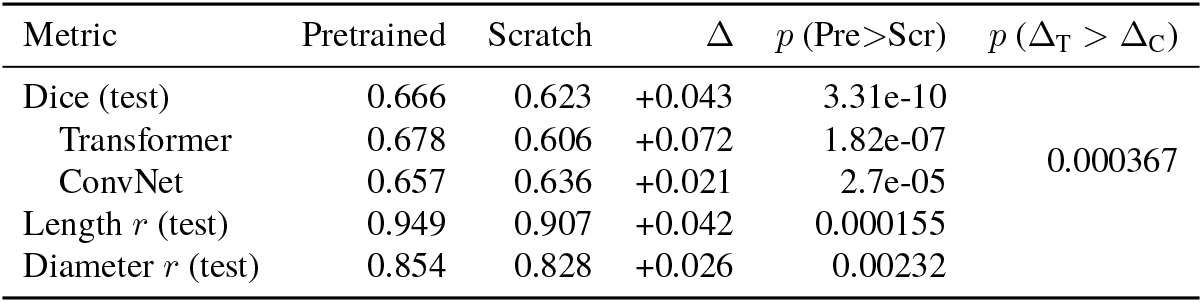
Summary of pretraining effects (Pretrained vs Scratch) across metrics. Architecture-specific breakdown shows Transformers benefit significantly more from pre-training than ConvNets; *p* (Δ_T_ *>* Δ_C_) tests whether this difference in benefit is significant.

**Table 5.** Validation and test performance (mean Dice) of each model, averaged across datasets; models are ranked by test Dice.

| Rank | Model | Arch | Val Dice | Test Dice | LR | Pre |
| --- | --- | --- | --- | --- | --- | --- |
| 1 | MobileSAM ViT-T | T | 0.7035 | 0.6925 | 1e-04 | ✓ |
| 2 | M2F Swin-S | T | 0.6985 | 0.6880 | 1e-04 | ✓ |
| 3 | M2F Swin-T | T | 0.7007 | 0.6847 | 1e-04 | ✓ |
| 4 | SegFormer B2 | T | 0.6977 | 0.6821 | 1e-04 |  |
| 5 | SegFormer B3 | T | 0.6942 | 0.6820 | 1e-04 |  |
| 6 | SegFormer B1 | T | 0.6912 | 0.6812 | 1e-04 | ✓ |
| 7 | M2F R50 | T | 0.6951 | 0.6769 | 1e-04 | ✓ |
| 8 | MA-Net Inc-v4 | C | 0.6869 | 0.6762 | 1e-04 | ✓ |
| 9 | U-Net++ Inc-v4 | C | 0.6825 | 0.6752 | 1e-04 | ✓ |
| 10 | U-Net++ R50 | C | 0.6879 | 0.6745 | 1e-04 | ✓ |
| 11 | LinkNet R50 | C | 0.6760 | 0.6689 | 1e-04 | ✓ |
| 12 | MA-Net R50 | C | 0.6884 | 0.6683 | 1e-04 | ✓ |
| 13 | DeepLabV3 R50 | C | 0.6837 | 0.6677 | 1e-04 |  |
| 14 | SAM2 Hiera-S | T | 0.6721 | 0.6643 | 1e-04 | ✓ |
| 15 | LinkNet Inc-v4 | C | 0.6783 | 0.6617 | 1e-03 | ✓ |
| 16 | SAM2 Hiera-B+ | T | 0.6661 | 0.6611 | 1e-04 | ✓ |
| 17 | DeepLabV3+ R50 | C | 0.6804 | 0.6534 | 1e-04 | ✓ |
| 18 | RootNav Hourglass | C | 0.6531 | 0.6441 | 1e-03 | ✓ |
| 19 | UNet-GN | C | 0.6536 | 0.6430 | 1e-04 | ✓ |
| 20 | UNet-GNRes | C | 0.6444 | 0.6351 | 1e-03 | ✓ |
| 21 | SegRoot W8xD5 | C | 0.6450 | 0.6344 | 1e-03 | ✓ |

**Figure 3.**
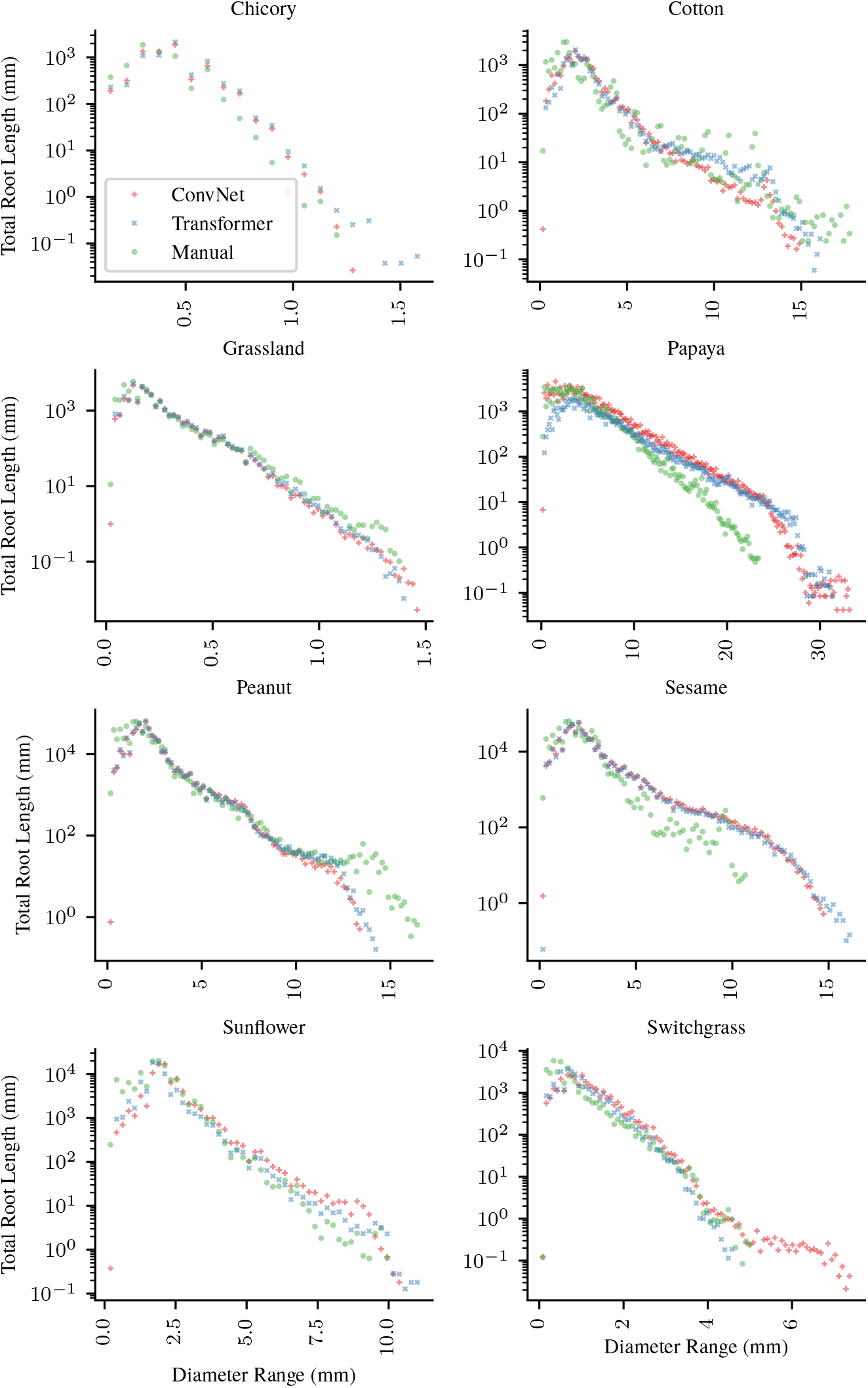
Diameter distribution per dataset. Each panel shows the percentage of total root length at each diameter bin (1 px resolution) for manual annotations, ConvNets (mean of top 4 from Table 5), and Transformers (mean of top 4 from Table 5). The log scale reveals the exponential decay of root length with increasing diameter.

The benefit of pre-training differed by architecture type. Transformer models had a mean Dice improvement of +0.072 from pre-training, compared to +0.022 for Con-vNets (*p* = 3.7 *×* 10^−4^; Table 4).

### Model rankings

Model rankings by test Dice are shown in Table 5. MobileSAM ViT-T had the highest test Dice (0.693), followed by M2F Swin-S (0.688) and M2F Swin-T (0.685). Rankings by root-length and root-diameter correlation are provided in Supplementary Tables 1 and 2. Some models ranked highly for Dice but lower for geometric agreement.

For example, MobileSAM led in Dice, but SegFormer B1 had the highest root-length correlation.

### Qualitative results

Figure 2 shows example segmentations from MobileSAM, the top-ranked model, across all nine datasets. Annotation errors are visible in the Sesame ground truth. In the Papaya example, the model captures root edges more accurately than the manual annotation.

### Dataset variation exceeds model variation

The variation in mean Dice scores across datasets (Table 7) was larger than the variation across model architectures (Table 5). Dataset choice explains 70.9% of variance in test Dice, far exceeding model choice (6.7%), pre-training (2.0%), architecture family (0.8%), learning rate (0.8%), and random seed (0.01%; Figure 8).

**Table 6.**
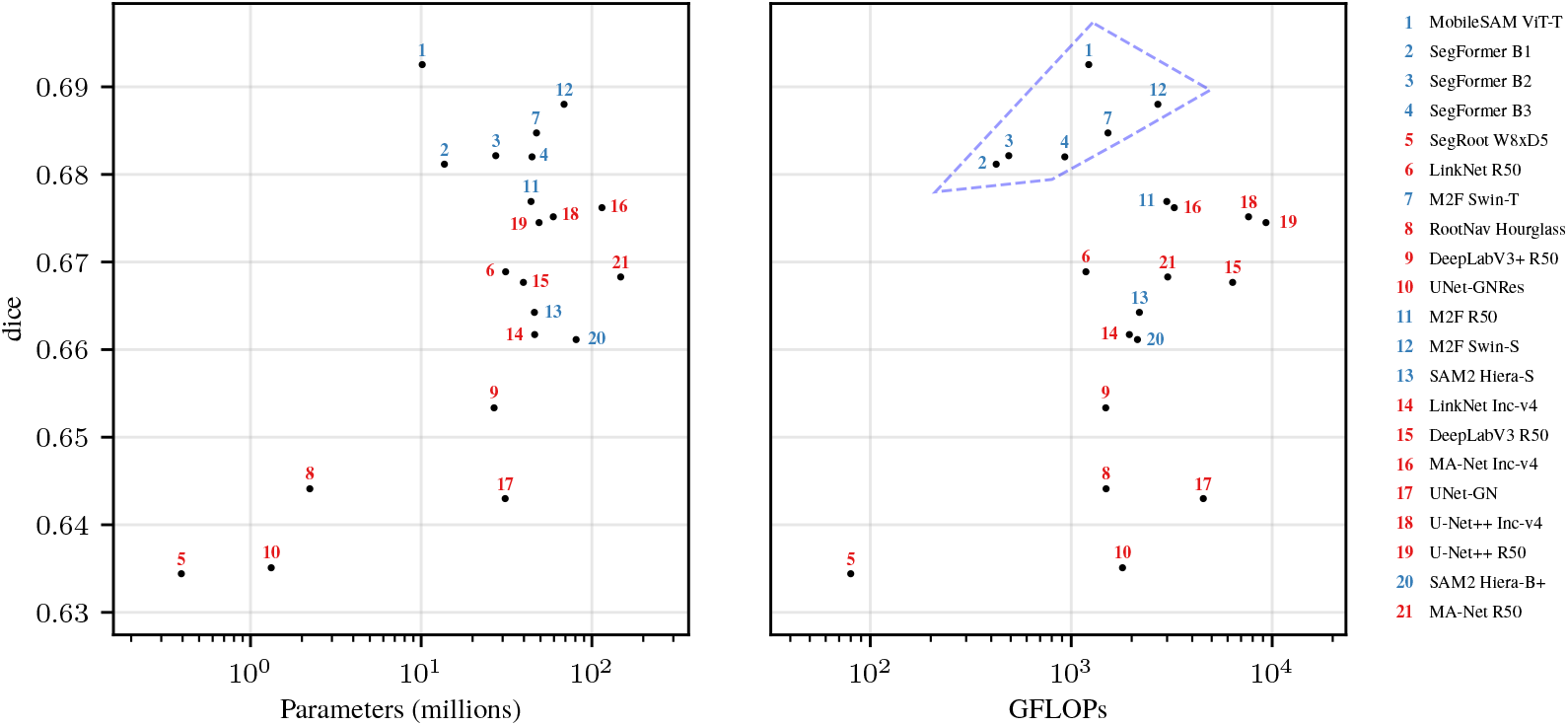
Test set dice vs model cost. Each model is assigned three ranks: dice rank (1 = highest dice), parameter rank (1 = fewest parameters), and FLOPs rank (1 = fewest FLOPs). The displayed number is the model’s position when sorted by the mean of these three ranks. Blue = Transformer, red = ConvNet. Dashed region highlights Transformers achieving higher Dice than ConvNets at similar or lower computational cost.

**Table 7.** Mean test performance per dataset, averaged across the selected models used in Table 5.

| Dataset | Mean Test Dice |
| --- | --- |
| Papaya | 0.8238 |
| Grassland | 0.7739 |
| Peanut | 0.7730 |
| Chicory | 0.6986 |
| Sunflower | 0.6957 |
| Switchgrass | 0.6911 |
| Cotton | 0.5965 |
| DeepRootLab | 0.4916 |
| Sesame | 0.4679 |

### Diameter distribution

The distribution of root length across diameter bins showed how models capture root morphology at different scales (Figures 3, 4b, 4a). For medium and large diameter roots, both ConvNet and Transformer architectures closely tracked manual annotations, following the characteristic exponential decay from the peak at approximately 5–10 pixels (Figure 4b). For the thinnest roots, however, models on average underestimated root length. At diameter bins below 5 pixels, manual annotations consistently attributed a higher percentage of total root length than either architecture (Figure 4a). At 2 pixels diameter, manual annotations showed 4.7% of root length compared to 1.4–1.7% for models; at 4 pixels, 8.2% compared to approximately 5%. Both ConvNet and Transformer models underestimated thin roots, either by failing to detect them or by predicting thicker diameters than annotated. This limitation was consistent across datasets (Figure 3) and is important for applications where accurate measurement of fine root structures is critical. At the thick end of the distribution, models predicted root length at diameter bins beyond the maximum annotated diameter, particularly in Papaya, Sesame, and Switchgrass (Figure 3). Inspection of the most extreme cases revealed that root merging contributes to this discrepancy: when adjacent roots run in parallel, models segment them as a single wide region, inflating the measured diameter (Figure 5).

**Figure 4.**
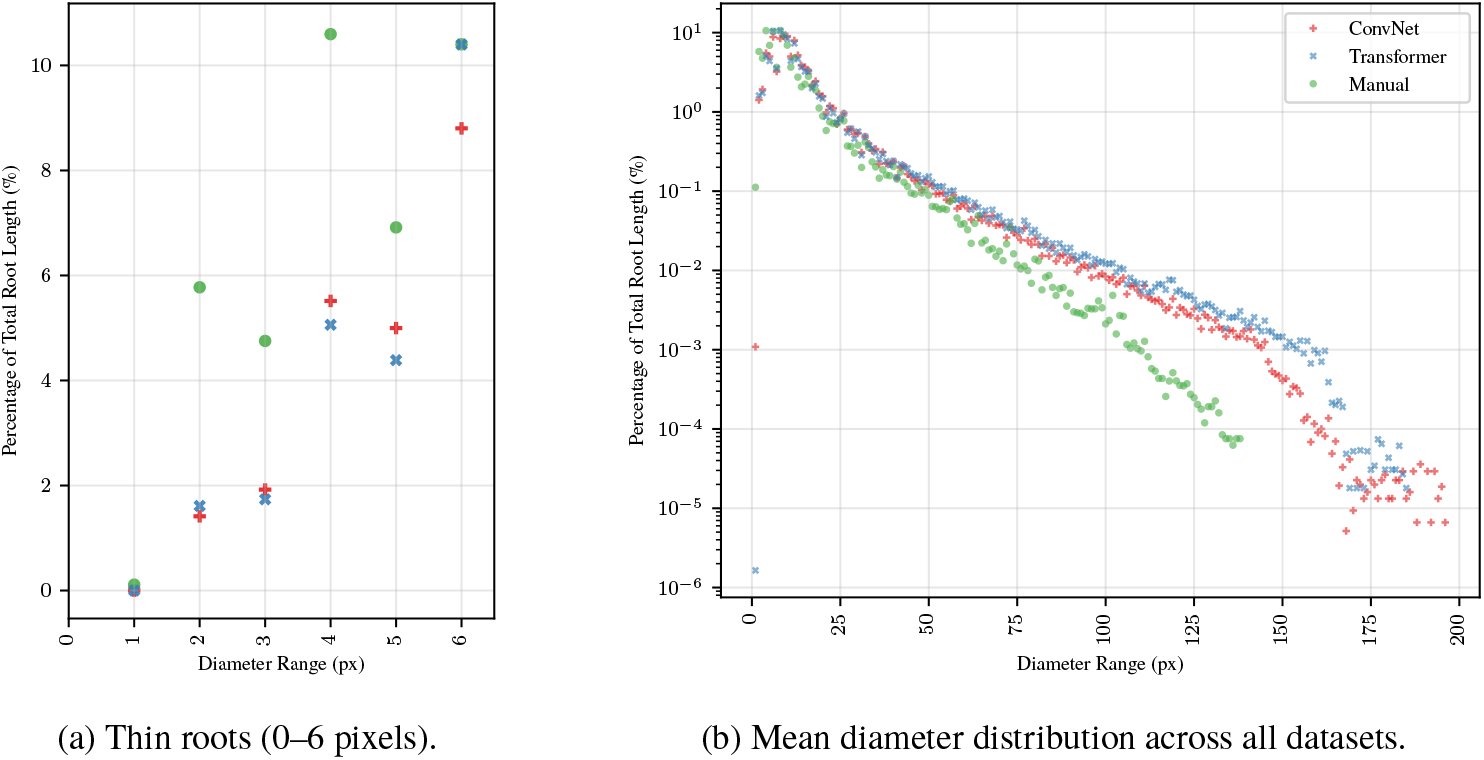
Diameter distributions averaged across datasets, showing manual annotations, ConvNets (mean of top 4 from Table 5), and Transformers (mean of top 4 from Table 5). (a) Thin roots showing underestimation by models on average. (b) Full range with log scale.

**Figure 5.**
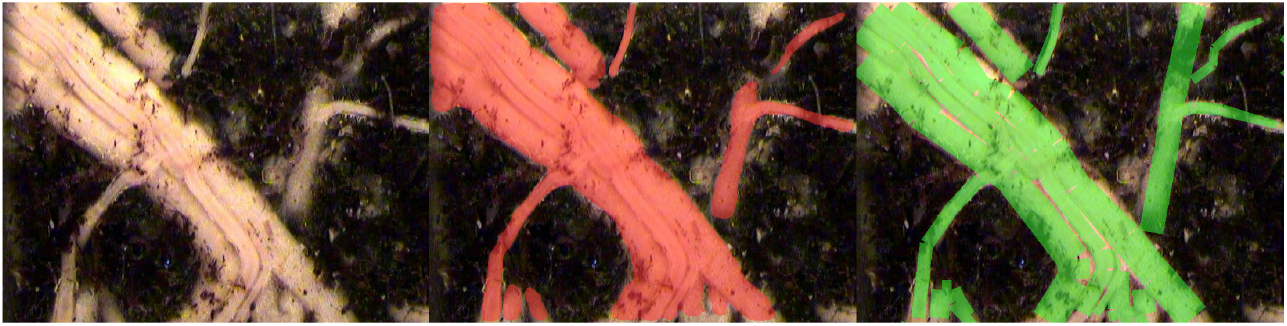
Root merging inflates diameter measurements. The image with the largest discrepancy between predicted and annotated maximum root diameter in the Papaya dataset is shown: original scan (left), MobileSAM prediction in red (center), and manual annotation in green (right). Adjacent roots that run in parallel are segmented as a single wide region by the model, causing RhizoVision Explorer to measure them as thick roots (predicted maximum diameter: 167 px) even though individual roots are thinner (annotated maximum diameter: 66 px). This root merging effect explains the model predictions at high diameter bins in Figure 3, particularly in Papaya, Sesame, and Switchgrass.

Thin root length underestimation fell into three dominant error types: RVE measurement artifacts from annotation corners (31%), roots missed entirely by the model (24%; Figure 6), and roots where the model segmentation was wider than the annotation (42%). Wider predictions occurred in all datasets, and missed roots in all datasets except Chicory. However, wider predictions do not always indicate a model error: in some cases the annotation was traced thinner than the actual root visible in the scan (Figure 7), indicating that annotation quality also contributes to thin root discrepancies.

## Discussion

### Transformer architectures outperform ConvNets

The consistent direction of Transformer improvement across Dice, root-length correlation, and root-diameter correlation supports H1, even though root-length correlation was not statistically significant (*p* = 0.099). This finding aligns with broader evidence from dense prediction tasks, where Vision Transformers produce more robust features than CNNs while relying less on texture cues [Jeeveswaran et al., 2022]. A systematic review of 36 medical imaging studies similarly found that Transformer-based models exhibit superior performance compared to CNNs, though with greater dependence on pre-training [Takahashi et al., 2024]. Our results extend these findings to root segmentation. One possible explanation is that the elongated, branching structure of roots benefits from Transformers’ self-attention mechanism, which integrates image-wide context from early layers in the network, unlike ConvNets which build up context gradually through successive layers. Other factors such as more effective transfer learning across large domain gaps may also contribute. The Transformer advantage was consistent across 8 of 9 datasets, with the largest improvement on DeepRootLab, suggesting the findings generalize to challenging field conditions.

### Pre-training benefits generalization

The consistent improvement from pre-training across all metrics shows that transfer learning provides useful initialisation despite the domain shift between pre-training datasets and root imagery, supporting H2. Transfer learning is widely adopted in agricultural imaging [Hossen et al., 2025], with reported benefits including faster convergence and reduced training time [Bosilj et al., 2020]. Our results confirm a modest but significant accuracy improvement (+0.043 Dice), though models trained from scratch still achieved reasonable performance.

Jeeveswaran et al. [2022] and Takahashi et al. [2024]claimed that Transformers are more data-hungry architectures, requiring larger datasets for effective training compared to ConvNets. The comparatively weak performance we observed for Transformers trained from scratch supports these claims, with a mean Dice of 0.606 compared to 0.636 for ConvNets trained from scratch. Zhou et al. [2021] hypothesised that pre-trained Transformers generalise better under large domain gaps, which may also contribute to the significantly greater benefit Transformers gained from pre-training compared to ConvNets (+0.072 vs +0.022 Dice; Table 4). Our pre-trained Transformer models used readily available weights from ImageNet, COCO, and Cityscapes, none of which contain root imagery, representing a large domain gap.

### Diameter distribution and thin root detection

Both Transformers and ConvNets underestimated thin roots, attributing less root length to the smallest diameter bins than manual annotations (Figure 4a). This parallels the “extremely small target” problem in medical image segmentation, where structures occupying few pixels can be ignored by networks trained with pixel-wise losses [Liu et al., 2021a]. Fine root structures also present known challenges across root phenotyping methods due to resolution limits [Weihs et al., 2024]. Our manual inspection of the largest thin root discrepancies showed that this reflects both genuine model limitations, roots missed entirely (Figure 6), and annotation inconsistency, where annotators traced roots thinner than they appear in the scan (Figure 7). The RVE measurement artifacts observed in 31% of inspected images arose from sharp corners in annotation masks produced by the annotation tool used for some datasets, and can likely be mitigated in practice using RVE’s built-in root pruning option, which removes short skeleton segments.

### Model efficiency and practical recommendations

For practitioners with constrained computational resources, pre-trained MobileSAM offers the best trade-off, achieving the highest Dice with fewer parameters than most ConvNets (Figure 6). More broadly, the highlighted region in Figure 6 shows that several Transformers (MobileSAM, SegFormer B1–B3, M2F Swin-T, M2F Swin-S) achieved high Dice with competitive FLOPs, forming an efficient frontier that no ConvNet matched. For applications where computational cost is the primary constraint and the segmentation task is relatively straightforward, SegRoot W8xD5 offers the lowest FLOPs of any evaluated model (Figure 6).

**Figure 6.**
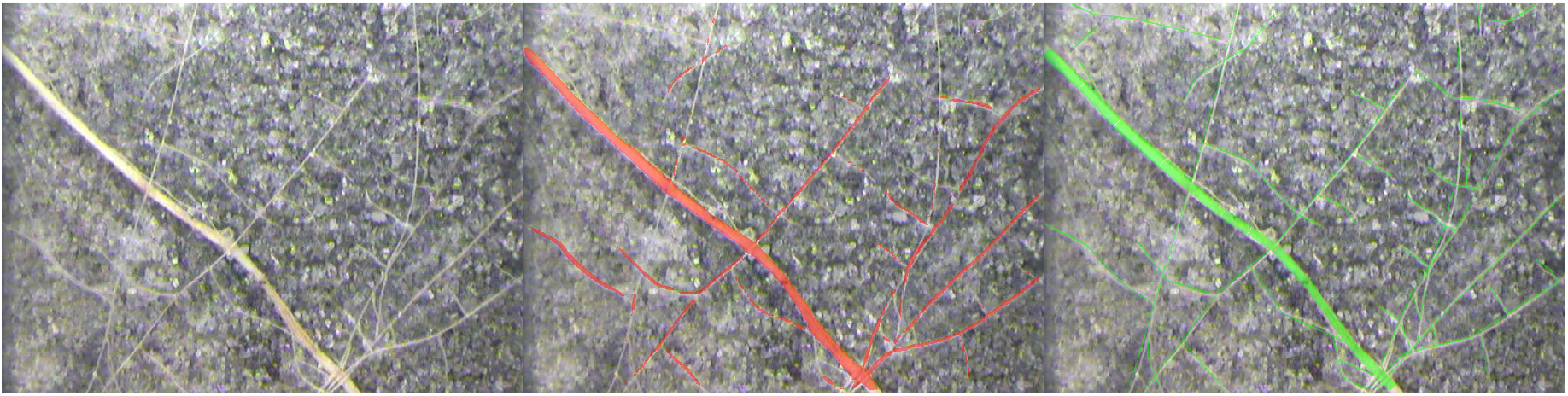
Thin roots missed by the model. The Peanut image with the largest thin-root length discrepancy (diameter bin 2, 2993 px) is shown: original scan (left), MobileSAM prediction in red (center), and manual annotation in green (right). Several fine lateral roots visible in the annotation are not captured by MobileSAM.

### Sources of variance

Dataset choice dominated performance variance (70.9%) compared to model choice (6.7%). Dataset-related factors, including inherent task difficulty, image quality, annotation quality, and dataset size, caused larger differences in segmentation performance than model architecture choice. Datasets can differ in inherent task difficulty due to factors such as root appearance, soil background complexity and artifacts, and species. The negligible effect of random seed (0.01%) shows high reproducibility. Given that dataset factors dominate performance variance, our results suggest practitioners should prioritize data curation over architecture selection. What constitutes a sufficient Dice for practical use will depend on the features of interest and dataset.

Annotation quality varied across datasets and likely contributed to the measured performance differences. In the Sesame dataset, we observed annotation errors including misaligned masks and incomplete root tracing (Figure 2), which contributed to lower Dice scores. For DeepRootLab, the corrective annotation approach likely contributes to lower Dice, as we only measure performance on harder regions of the image where models are more likely to have errors. Because these annotations were created by correcting errors, they may disproportionately capture the differences between stronger and weaker models, which could explain why DeepRootLab showed the widest spread in per-model Dice (range 0.33 vs ≤ 0.08 for all other datasets; Supplementary Figures 1–2). Despite this, the overall model ranking was stable: removing DeepRootLab and averaging the remaining eight datasets yielded a ranking highly correlated with the full nine-dataset ranking (Spearman *ρ* = 0.92). Additionally, this dataset spans eleven species imaged over multiple years under field conditions, introducing challenging variation in root appearance and age.

Manual inspection of thin root discrepancies further showed that some cases where the model predicted thicker roots were in fact annotation errors, where roots were traced thinner than they appear in the scan (Figure 7). These annotation issues highlight a broader challenge: when models segment roots more accurately than annotators, standard metrics penalize the model.

**Figure 7.**
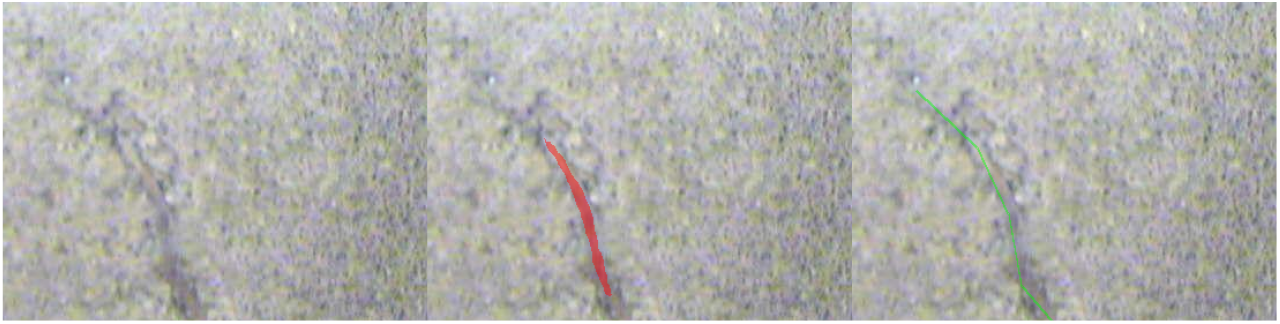
Annotation error in Cotton: roots traced too thin. Top half of the Cotton image with the largest “predicted too thick” discrepancy in diameter bin 2 ( − 353 px): original scan (left), MobileSAM prediction in red (center), and manual annotation in green (right). The model under-segments some root length but captures root thickness more accurately than the annotation, which traces roots thinner than they appear in the scan.

**Figure 8.**
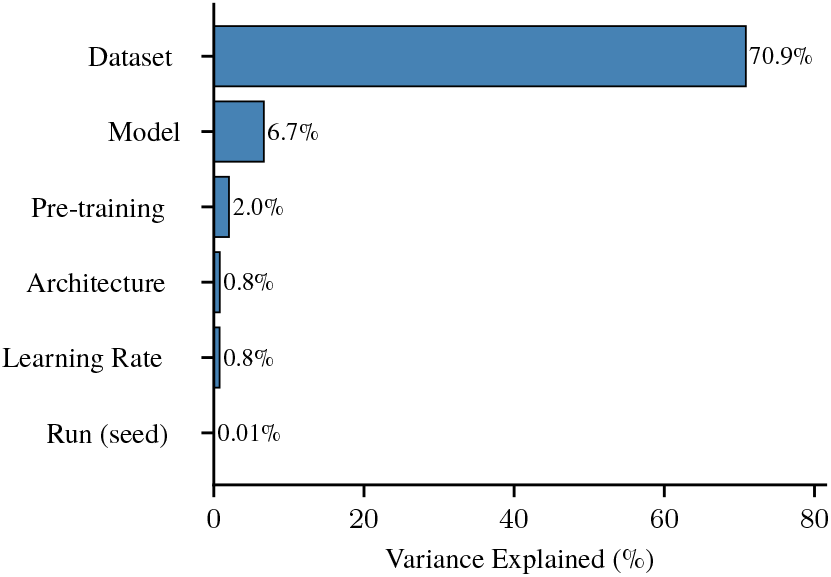
Proportion of variance in test Dice (*η*^2^)attributable to each experimental factor. Dataset choice alone accounts for 70.9% of performance variance, farexceeding model architecture, pre-training strategy, or hyperparameter selection.

### Limitations and future work

Pre-trained weights were sourced from ImageNet, COCO, Cityscapes, or model-specific datasets, reflecting the practical reality that researchers typically fine-tune from publicly available checkpoints. The consistent advantage of pre-training across this heterogeneous set of sources shows that H2 holds regardless of the specific pre-training dataset used. Domain-specific pre-training on root imagery could yield further gains. The discrepancy between model and annotation at thin root diameters (Figure 4a), which reflects both genuine model limitations and annotation inconsistencies (Figures 6 and 7), indicates that improvements in both segmentation models and annotation protocols would benefit root phenotyping applications.

## Conclusion

We evaluated 21 deep learning architectures across nine root image datasets (1,511 training runs). Transformer architectures outperformed ConvNets (*p* = 1.5 × 10^−3^), with MobileSAM achieving the highest Dice (0.693). Pre-training improved performance (*p* = 3.3 × 10^−10^), particularly for Transformers (+0.072 vs +0.022 Dice, *p* = 3.7 × 10^−4^). Dataset choice explained 70.9% of performance variance, far exceeding model choice (6.7%). For practitioners, investing in training data quality and quantity matters more than architecture selection. Among models, pre-trained MobileSAM offers strong accuracy with low computational cost.

