## Supplementary Materials for "Transformers Outperform ConvNets for Root Segmentation: A Systematic Comparison Across Nine Datasets"

##### Supplementary Table 1: Model ranking by root-length correlation

Root length agreement (Pearson correlation $r$) on the test set, averaged across datasets; configurations are selected using validation Dice, and models are ranked by test $r$.

| Rank | Model | Arch | Length $r$ (test) | Pre |
|---|---|---|---|---|
| 1 | SegFormer B1 | T | 0.958 | ✓ |
| 2 | MA-Net Inc-v4 | C | 0.958 | ✓ |
| 3 | SegFormer B3 | T | 0.956 | |
| 4 | MA-Net R50 | C | 0.955 | ✓ |
| 5 | LinkNet Inc-v4 | C | 0.955 | ✓ |
| 6 | M2F Swin-T | T | 0.955 | ✓ |
| 7 | U-Net++ Inc-v4 | C | 0.954 | ✓ |
| 8 | SegFormer B2 | T | 0.953 | |
| 9 | M2F Swin-S | T | 0.952 | ✓ |
| 10 | M2F R50 | T | 0.952 | ✓ |
| 11 | LinkNet R50 | C | 0.952 | ✓ |
| 12 | MobileSAM ViT-T | T | 0.950 | ✓ |
| 13 | DeepLabV3 R50 | C | 0.950 | |
| 14 | U-Net++ R50 | C | 0.950 | ✓ |
| 15 | SAM2 Hiera-B+ | T | 0.949 | ✓ |
| 16 | DeepLabV3+ R50 | C | 0.948 | ✓ |
| 17 | SegRoot W8xD5 | C | 0.944 | ✓ |
| 18 | RootNav Hourglass | C | 0.943 | ✓ |
| 19 | SAM2 Hiera-S | T | 0.943 | ✓ |
| 20 | UNet-GNRes | C | 0.934 | ✓ |
| 21 | UNet-GN | C | 0.933 | ✓ |

##### Supplementary Table 2: Model ranking by root-diameter correlation

Root diameter agreement (Pearson correlation $r$) on the test set, averaged across datasets; configurations are selected using validation Dice, and models are ranked by test $r$.

| Rank | Model | Arch | Diameter $r$ (test) | Pre |
|---|---|---|---|---|
| 1 | M2F Swin-S | T | 0.875 | ✓ |
| 2 | SegFormer B3 | T | 0.874 | |
| 3 | MobileSAM ViT-T | T | 0.872 | ✓ |
| 4 | SegFormer B1 | T | 0.871 | ✓ |
| 5 | SegFormer B2 | T | 0.869 | |
| 6 | M2F R50 | T | 0.865 | ✓ |
| 7 | LinkNet Inc-v4 | C | 0.864 | ✓ |
| 8 | SAM2 Hiera-B+ | T | 0.861 | ✓ |
| 9 | U-Net++ R50 | C | 0.858 | ✓ |
| 10 | MA-Net R50 | C | 0.857 | ✓ |
| 11 | MA-Net Inc-v4 | C | 0.853 | ✓ |
| 12 | U-Net++ Inc-v4 | C | 0.851 | ✓ |
| 13 | SAM2 Hiera-S | T | 0.850 | ✓ |
| 14 | UNet-GN | C | 0.848 | ✓ |
| 15 | UNet-GNRes | C | 0.848 | ✓ |
| 16 | LinkNet R50 | C | 0.847 | ✓ |
| 17 | DeepLabV3+ R50 | C | 0.845 | ✓ |
| 18 | SegRoot W8xD5 | C | 0.841 | ✓ |
| 19 | DeepLabV3 R50 | C | 0.838 | |
| 20 | RootNav Hourglass | C | 0.822 | ✓ |
| 21 | M2F Swin-T | T | 0.815 | ✓ |

**Supplementary Table 3: Dataset splits**

Number of images in each dataset split, with image resolution and DPI.

| Dataset | Train | Val | Test | Total | DPI | px/mm | Image Size (px) |
|---|---|---|---|---|---|---|---|
| Chicory | 29 | 9 | 10 | 48 | 338 | 13.3 | 3991×1842 |
| Cotton | 1,271 | 564 | 577 | 2,412 | 150 | 5.9 | 736×552 |
| DeepRootLab | 262 | 86 | 90 | 438 | 1300 | 51.2 | 1224×1024 |
| Grassland | 71 | 21 | 16 | 108 | 1200 | 47.2 | 2550×2196 |
| Papaya | 282 | 131 | 133 | 546 | 150 | 5.9 | 736×552 |
| Peanut | 11,485 | 3,347 | 4,793 | 19,625 | 150 | 5.9 | 736×552 |
| Sesame | 8,637 | 2,625 | 3,084 | 14,346 | 150 | 5.9 | 736×552 |
| Sunflower | 2,211 | 722 | 967 | 3,900 | 120 | 4.7 | 640×480 |
| Switchgrass | 2,647 | 665 | 600 | 3,912 | 300 | 11.8 | 510×720 |
| **Total** | **26,895** | **8,170** | **10,270** | **45,335** | | | |

**Supplementary Table 4: Per-dataset test Dice (pre-trained)**

Test Dice for each pre-trained model on each dataset. For each model, the best learning rate is selected by mean validation Dice across datasets. Models are ranked by mean test Dice (descending). T = Transformer, C = ConvNet.

| Model | Type | LR | Chicory | Cotton | DeepRootLab | Grassland | Papaya | Peanut | Sesame | Sunflower | Switchgrass |
|---|---|---|---|---|---|---|---|---|---|---|---|
| MobileSAM ViT-T | T | $10^{-4}$ | 0.705 | 0.626 | 0.639 | 0.789 | 0.833 | 0.784 | 0.469 | 0.706 | 0.682 |
| M2F Swin-S | T | $10^{-4}$ | 0.705 | 0.620 | 0.596 | 0.782 | 0.832 | 0.781 | 0.467 | 0.684 | 0.726 |
| M2F Swin-T | T | $10^{-4}$ | 0.706 | 0.615 | 0.569 | 0.792 | 0.837 | 0.778 | 0.474 | 0.675 | 0.716 |
| SegFormer B1 | T | $10^{-4}$ | 0.692 | 0.627 | 0.529 | 0.777 | 0.832 | 0.778 | 0.483 | 0.699 | 0.715 |
| SegFormer B3 | T | $10^{-4}$ | 0.693 | 0.616 | 0.545 | 0.768 | 0.838 | 0.786 | 0.464 | 0.675 | 0.725 |
| M2F R50 | T | $10^{-4}$ | 0.707 | 0.607 | 0.546 | 0.772 | 0.831 | 0.772 | 0.467 | 0.693 | 0.699 |
| MA-Net Inc-v4 | C | $10^{-4}$ | 0.708 | 0.604 | 0.532 | 0.781 | 0.826 | 0.784 | 0.462 | 0.707 | 0.683 |
| U-Net++ Inc-v4 | C | $10^{-4}$ | 0.708 | 0.593 | 0.509 | 0.781 | 0.827 | 0.782 | 0.470 | 0.715 | 0.691 |
| U-Net++ R50 | C | $10^{-4}$ | 0.708 | 0.603 | 0.503 | 0.780 | 0.830 | 0.777 | 0.472 | 0.707 | 0.690 |
| SegFormer B2 | T | $10^{-4}$ | 0.692 | 0.605 | 0.476 | 0.779 | 0.834 | 0.785 | 0.467 | 0.703 | 0.725 |
| LinkNet R50 | C | $10^{-4}$ | 0.698 | 0.586 | 0.509 | 0.771 | 0.825 | 0.776 | 0.467 | 0.703 | 0.685 |
| MA-Net R50 | C | $10^{-4}$ | 0.705 | 0.599 | 0.480 | 0.782 | 0.816 | 0.769 | 0.470 | 0.713 | 0.681 |
| SAM2 Hiera-S | T | $10^{-4}$ | 0.704 | 0.570 | 0.459 | 0.781 | 0.831 | 0.773 | 0.461 | 0.706 | 0.692 |
| LinkNet Inc-v4 | C | $10^{-3}$ | 0.703 | 0.600 | 0.414 | 0.786 | 0.823 | 0.768 | 0.470 | 0.712 | 0.678 |
| SAM2 Hiera-B+ | T | $10^{-4}$ | 0.701 | 0.581 | 0.442 | 0.776 | 0.830 | 0.772 | 0.475 | 0.682 | 0.692 |
| DeepLabV3+ R50 | C | $10^{-4}$ | 0.687 | 0.598 | 0.450 | 0.770 | 0.825 | 0.774 | 0.453 | 0.657 | 0.667 |
| DeepLabV3 R50 | C | $10^{-4}$ | 0.664 | 0.598 | 0.454 | 0.756 | 0.823 | 0.774 | 0.448 | 0.685 | 0.667 |
| RootNav Hourglass | C | $10^{-3}$ | 0.700 | 0.545 | 0.401 | 0.762 | 0.809 | 0.743 | 0.466 | 0.686 | 0.684 |
| UNet-GN | C | $10^{-4}$ | – | 0.577 | 0.403 | 0.749 | 0.797 | 0.770 | 0.464 | 0.690 | 0.692 |
| UNet-GNRes | C | $10^{-3}$ | – | 0.575 | 0.307 | 0.782 | 0.810 | 0.764 | 0.470 | 0.700 | 0.673 |
| SegRoot W8xD5 | C | $10^{-3}$ | 0.681 | 0.560 | 0.367 | 0.734 | 0.801 | 0.738 | 0.466 | 0.683 | 0.680 |

##### Supplementary Table 5: Per-dataset test Dice (trained from scratch)

Test Dice for each model trained from scratch on each dataset. For each model, the best learning rate is selected by mean validation Dice across datasets. Models are ranked by mean test Dice (descending). T = Transformer, C = ConvNet.

| Model | Type | LR | Chicory | Cotton | DeepRootLab | Grassland | Papaya | Peanut | Sesame | Sunflower | Switchgrass |
|---|---|---|---|---|---|---|---|---|---|---|---|
| SegFormer B2 | T | $10^{-4}$ | 0.687 | 0.619 | 0.563 | 0.770 | 0.835 | 0.785 | 0.469 | 0.695 | 0.716 |
| SegFormer B3 | T | $10^{-4}$ | 0.693 | 0.628 | 0.546 | 0.777 | 0.825 | 0.786 | 0.469 | 0.705 | 0.708 |
| SegFormer B1 | T | $10^{-4}$ | 0.686 | 0.620 | 0.501 | 0.773 | 0.827 | 0.781 | 0.475 | 0.663 | 0.716 |
| DeepLabV3 R50 | C | $10^{-4}$ | 0.674 | 0.595 | 0.561 | 0.757 | 0.825 | 0.777 | 0.463 | 0.692 | 0.664 |
| LinkNet Inc-v4 | C | $10^{-3}$ | 0.703 | 0.578 | 0.385 | 0.778 | 0.814 | 0.758 | 0.466 | 0.682 | 0.685 |
| RootNav Hourglass | C | $10^{-4}$ | 0.701 | 0.546 | 0.378 | 0.767 | 0.804 | 0.751 | 0.466 | 0.697 | 0.686 |
| U-Net++ R50 | C | $10^{-4}$ | 0.701 | 0.562 | 0.343 | 0.762 | 0.794 | 0.761 | 0.471 | 0.701 | 0.682 |
| MA-Net R50 | C | $10^{-4}$ | 0.694 | 0.566 | 0.352 | 0.756 | 0.803 | 0.752 | 0.472 | 0.700 | 0.678 |
| U-Net++ Inc-v4 | C | $10^{-4}$ | 0.700 | 0.587 | 0.273 | 0.763 | 0.814 | 0.768 | 0.471 | 0.693 | 0.679 |
| MA-Net Inc-v4 | C | $10^{-4}$ | 0.699 | 0.567 | 0.271 | 0.768 | 0.816 | 0.760 | 0.469 | 0.695 | 0.677 |
| DeepLabV3+ R50 | C | $10^{-4}$ | 0.683 | 0.584 | 0.342 | 0.733 | 0.799 | 0.757 | 0.467 | 0.670 | 0.675 |
| LinkNet R50 | C | $10^{-3}$ | 0.702 | 0.480 | 0.331 | 0.768 | 0.814 | 0.748 | 0.467 | 0.700 | 0.672 |
| M2F R50 | T | $10^{-4}$ | 0.691 | 0.560 | 0.292 | 0.747 | 0.801 | 0.753 | 0.469 | 0.691 | 0.670 |
| MobileSAM ViT-T | T | $10^{-4}$ | 0.666 | 0.564 | 0.277 | 0.731 | 0.800 | 0.754 | 0.467 | 0.692 | 0.684 |
| UNet-GNRes | C | $10^{-3}$ | 0.703 | 0.567 | 0.169 | 0.771 | 0.811 | 0.766 | 0.471 | 0.698 | 0.669 |
| UNet-GN | C | $10^{-4}$ | 0.702 | 0.555 | 0.162 | 0.767 | 0.803 | 0.768 | 0.460 | 0.695 | 0.690 |
| M2F Swin-S | T | $10^{-4}$ | 0.699 | 0.552 | 0.178 | 0.760 | 0.792 | 0.763 | 0.464 | 0.687 | 0.672 |
| M2F Swin-T | T | $10^{-4}$ | 0.701 | 0.553 | 0.091 | 0.761 | 0.804 | 0.764 | 0.468 | 0.684 | 0.700 |
| SegRoot W8xD5 | C | $10^{-3}$ | 0.685 | 0.553 | 0.086 | 0.735 | 0.803 | 0.730 | 0.455 | 0.685 | 0.664 |
| SAM2 Hiera-B+ | T | $10^{-4}$ | 0.660 | 0.122 | 0.196 | 0.716 | 0.581 | 0.652 | 0.380 | 0.592 | 0.617 |
| SAM2 Hiera-S | T | $10^{-3}$ | 0.118 | 0.200 | 0.182 | 0.532 | 0.715 | 0.598 | 0.374 | 0.519 | 0.576 |

##### Supplementary Figure 1: Mean test Dice across datasets

Mean test Dice for each model across all nine datasets, using the best configuration selected by mean validation Dice. Models ordered by mean test Dice (best at left). Markers: × = Transformer, + = ConvNet.

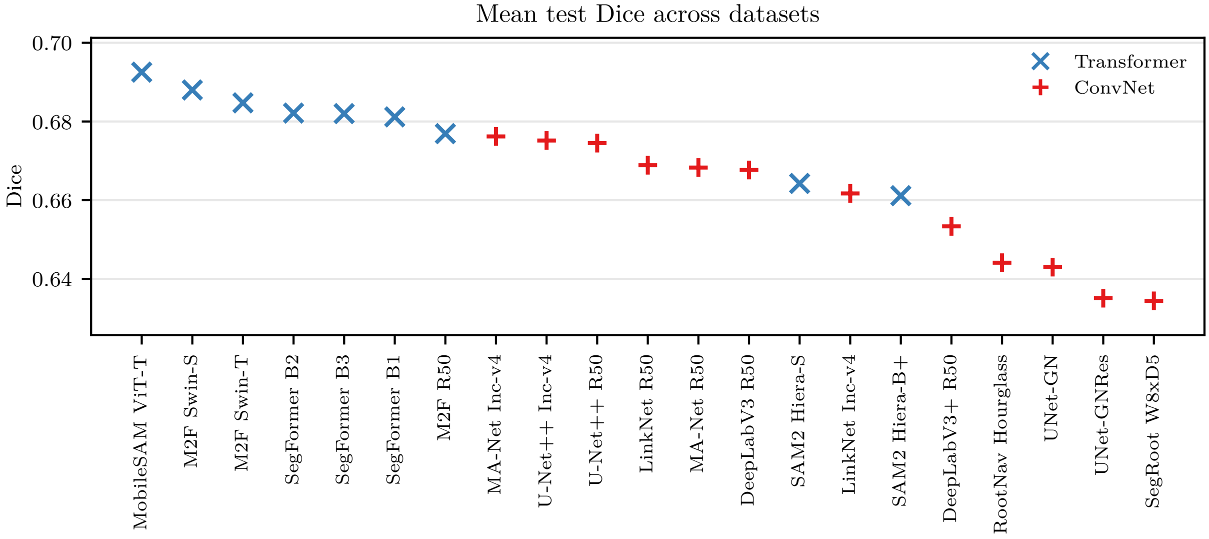


**Supplementary Figure 2: Per-dataset test Dice breakdown**

Test Dice for each model on each of the nine datasets. Same model ordering and configuration selection as Supplementary Figure 1. Markers: × = Transformer, + = ConvNet.

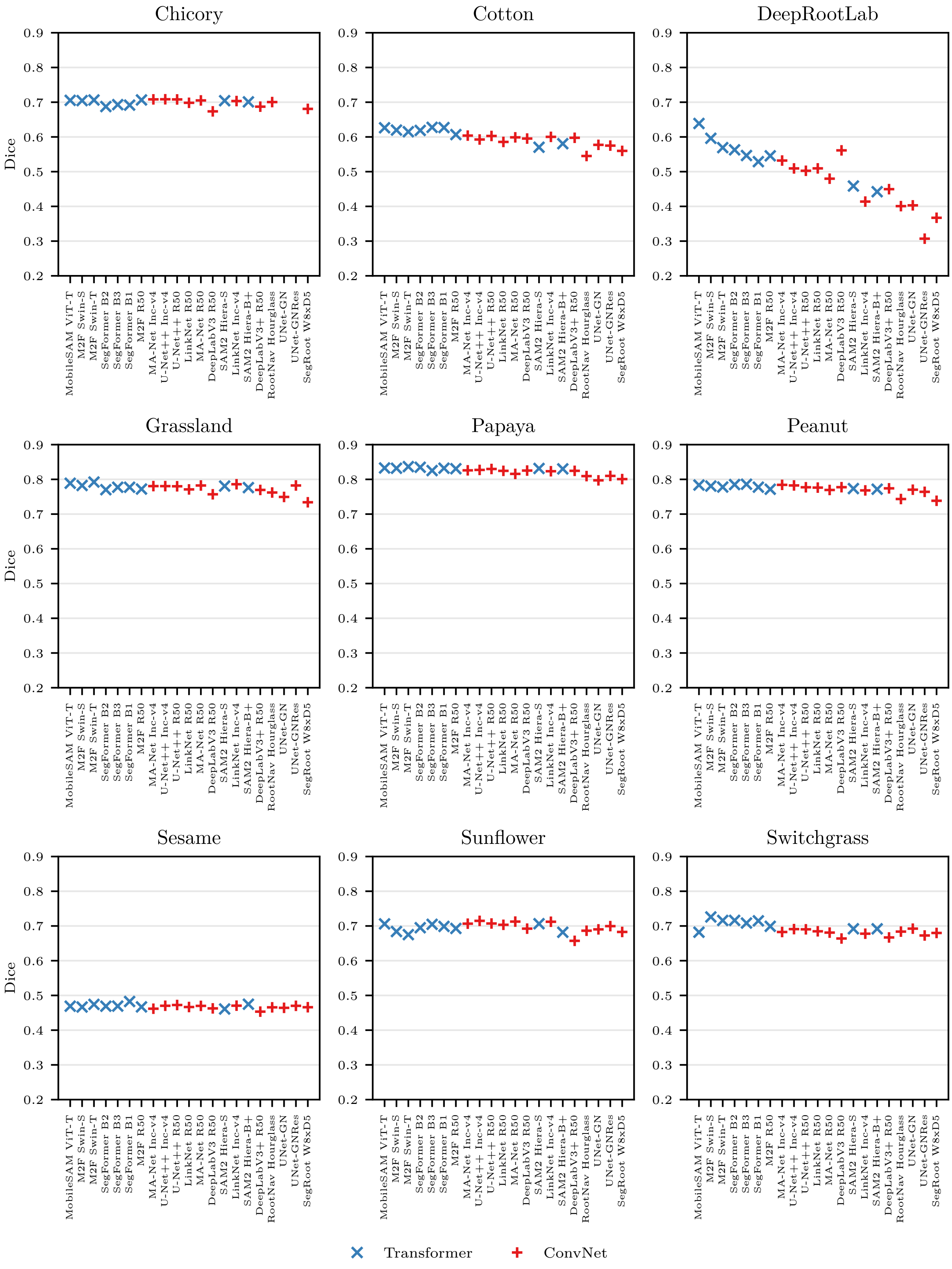
Chicory
Cotton
DeepRootLab
Grassland
Papaya
Peanut
Sesame
Sunflower
Switchgrass
Dice
0.9
0.8
0.7
0.6
0.5
0.4
0.3
0.2
MobileSAM ViT-T
M2F Swin-S
M2F Swin-T
SegFormer B2
SegFormer B3
SegFormer B1
M2F R50
MA-Net Inc-v4
U-Net++ Inc-v4
U-Net++ R50
LinkNet R50
MA-Net R50
DeepLabV3 R50
SAM2 Hiera-S
LinkNet Inc-v4
SAM2 Hiera-B+
DeepLabV3+ R50
RootNav Hourglass
UNet-GN
UNet-GNRes
SegRoot W8xD5
Transformer
ConvNet

### RhizoVision Explorer Configuration

Root phenotyping measurements were extracted using a fork of RhizoVision Explorer (RVE) v2.0 with patches for headless operation, memory safety, and command-line argument parsing. The fork and its modified dependency library are available at `https://github.com/sotlampr/RhizoVisionExplorer` (commit `a08860c`) and `https://github.com/sotlampr/cvutil` (commit `160d38c`).

RVE was invoked as: `rv -i -r -na -dranges 1,2,3,...,201`, where `-i` inverts the image (white roots on black background), `-r` enables recursive directory processing, `-na` prevents appending to existing output files, and `-dranges` specifies diameter bin boundaries from 1 to 201 pixels in 1-pixel increments.
